# Swan: a library for the analysis and visualization of long-read transcriptomes

**DOI:** 10.1101/2020.06.09.143024

**Authors:** Fairlie Reese, Ali Mortazavi

## Abstract

**Motivation:** Long-read RNA-sequencing technologies such as PacBio and Oxford Nanopore have discovered an explosion of new transcript isoforms that are difficult to visually analyze using currently available tools. We introduce the Swan Python library, which is designed to analyze and visualize transcript models.

**Results:** Swan finds 4,909 differentially expressed transcripts between cell lines HepG2 and HFFc6, including 279 that are differentially expressed even though the parent gene is not. Additionally, Swan discovers 1,021 reproducible exon skipping and 73 intron retention events not recorded in the GENCODE v29 annotation.

**Availability:** The Swan library for Python 3 is available on PyPi and on GitHub at https://pypi.org/project/swan-vis/1.0/ and https://github.com/mortazavilab/swan_paper.

## Introduction

Alternative splicing plays a critical role in many biological processes and disease states. However, standard genomic assays have difficulties capturing the comprehensive spectrum of full-length alternative splicing due to the limitations of short reads in reconstructing transcript isoforms (Conesa et al., 2016). Long-read sequencing platforms such as PacBio and Oxford Nanopore have led to an explosion in discovery of transcript isoforms that were impossible to assemble with short reads. Current transcript model visualization tools such as Sashimi or LeafCutter plots are primarily designed for visualizing short reads rather than leveraging full-length isoforms (Katz et al., 2015; Li et al., 2018). Furthermore, these tools display isoforms on a genomic scale which complicates interpreting and distinguishing similar isoforms from one another.

We introduce the Swan Python library, which is designed to analyze and visualize transcript models. Swan’s graphical model approach allows the user to visually distinguish between transcript models and to identify novel exon skipping and intron retention events commonly missed by short reads. Furthermore, Swan incorporates statistical models to detect differential gene and transcript expression, enabling quantitative comparison of full-length transcript models in different biological settings. We demonstrate Swan’s utility by applying it to full-length PacBio transcriptome data from the HepG2 and HFFc6 human cell lines, which are publicly available on the ENCODE portal.

## Methods

### 2.1 Input and transcriptome representation

Swan works by processing transcript models from GTF files or from a TALON database (Wyman et al., 2019) into a SwanGraph data structure consisting of a series of data frames and a graph. Each unique genomic coordinate, exon, intron, and transcript is recorded in the data frames and used to construct a graphical representation of the transcriptome. Genomic locations are represented as nodes. Introns and exons are represented as edges between nodes, and a full transcript is represented as a path that traverses the nodes and edges present in the transcript. This flexible scheme makes it possible to add additional datasets and track which nodes, edges, and full transcripts are present in each. Expression data can also be added from a tab-delimited counts matrix indexed by the transcript ID with columns for each dataset and is tracked alongside each transcript’s presence in each dataset.

### 2.2 Visualization and analysis

Swan plots directly correspond to the graphical representation in the SwanGraph. Nodes and edges are colored according to their role in the gene or transcript **(Figure 1)**. Transcription start site (TSSs) nodes are colored blue, transcription end site (TESs) nodes are colored orange, and internal nodes that are traversed between the TSS and TES of a transcript are colored yellow. Nodes with more than one role such as a node that can be either a TSS or an internal node are preferentially colored by their most “unique” role in the gene (TES > TSS > internal). Exonic edges are colored green and intronic edges are colored pink. Nodes are spaced out evenly regardless of genomic location. This facilitates visual recognition of splicing events such as alternative 5’/3’ splice site usage, which sometimes occur only a few base pairs away from the canonical splice site. Such events are difficult to distinguish on a genomic scale as the difference between splice sites in a genome browser-style representation consists of a difference of just a few pixels. In contrast, Swan’s visualization uses an entirely different node to draw attention to splicing complexity.

**Figure 1:**
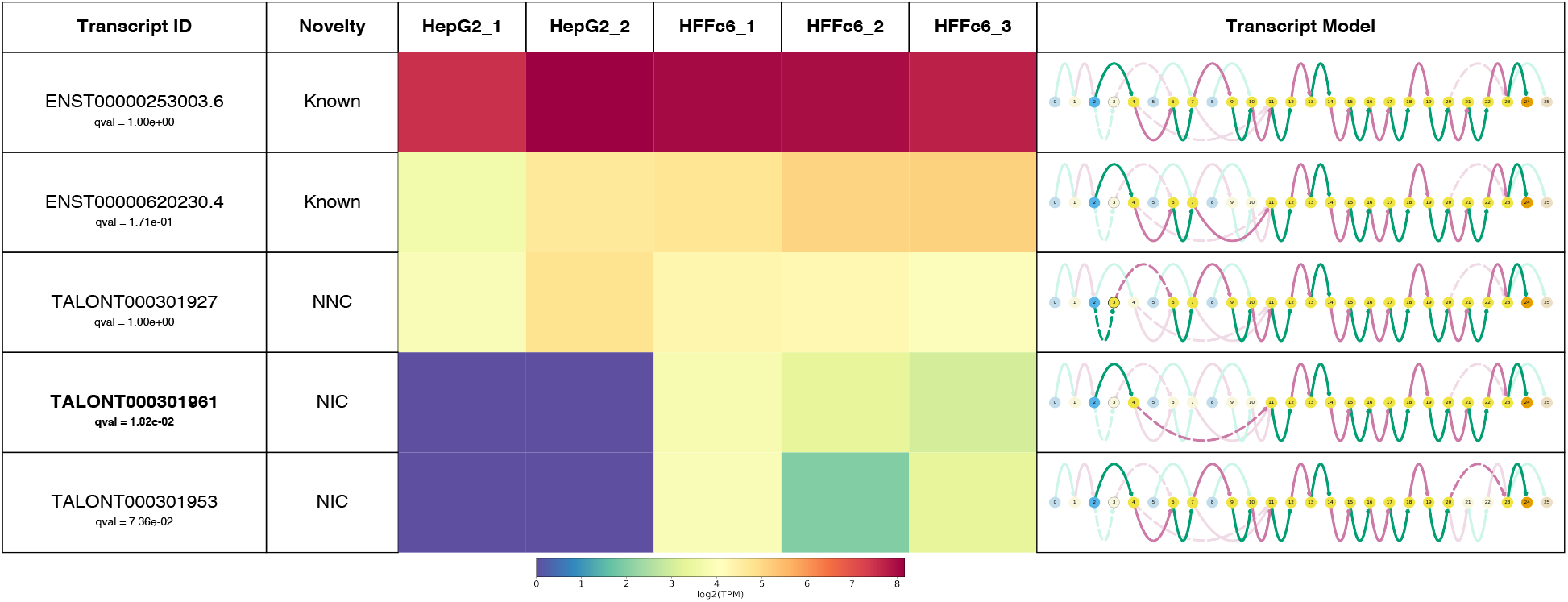
Swan report for expressed *Adrm1* transcripts. Bolded entries are differentially expressed transcripts between human cell lines HepG2 and HFFc6 (q <= 0.05). Transcript novelty categories are determined by TALON. Novel splice sites are outlined nodes and novel combinations of splice sites are dashed edges. TSS nodes are blue and TES nodes are orange. Exonic edges are green and intronic edges are pink.

Gene summary graphs provide an overview of the splicing complexity of a specific gene. Transcript path graphs show the nodes and edges traversed by a specific transcript model in bright colors through the rest of the gene graph, where unused nodes and edges are grayed out. In both graph plots, there is an option to draw attention to nodes/edges that are not present in the annotation or to nodes/edges that are in a specific dataset. Transcript paths can also be plotted using a traditional genome browser-style representation to provide a familiar reference.

Swan generates gene-level PDF reports showing the transcript path representation for each transcript model for a given gene, along with the presence or absence of each isoform in the datasets included. Swan also offers several analysis tools to supplement its visualization suite. When transcript abundance information is provided, differential expression tests for genes and transcripts, implemented via Scanpy’s diffxpy, can be run and visualized in Swan reports (Wolf et al., 2018). Swan also detects novel exon skipping and intron retention events.

## Results

We obtained mapped PacBio RNA-seq datasets from the ENCODE portal for 2 replicates of HepG2 and 3 replicates of HFFc6. We used TALON v5 to call transcript isoforms and filter novel ones for high reproducibility for 22,857 transcript models in HepG2 and 28,814 in HFFc6. We then used TALON to obtain GTFs and abundance files for each dataset. We fed the resulting transcript models and abundance information into Swan, along with the GENCODE v29 annotation (Harrow et al., 2012).

Swan identified 4,009 differentially expressed genes and 4,909 differentially expressed transcripts across cell lines. Of the differentially expressed transcripts, 279 belong to a parent gene that is not differentially expressed and are therefore candidates for isoform switching analysis. Additionally, Swan found 285 novel exon skipping events and 73 novel intron retention events.

Swan provides a platform for deeply exploring full-length transcriptome data. Its intuitive visualizations and flexible analysis tools enable discovery of novel splicing events and differential transcript expression. In the future, we hope to expand Swan’s capabilities in many ways including implementing multi-way differential expression tests and adapting it to work with single cell long-read RNA-seq datasets.

